# Data-driven algorithm for the diagnosis of behavioral variant frontotemporal dementia

**DOI:** 10.1101/2019.12.19.883462

**Authors:** Ana L. Manera, Mahsa Dadar, John van Swieten, Barbara Borroni, Raquel Sanchez-Valle, Fermin Moreno, Robert LaForce, Caroline Graff, Matthis Synofzik, Daniela Galimberti, James Rowe, Mario Masellis, Maria Carmela Tartaglia, Elizabeth Finger, Rik Vandenberghe, Alexandre de Mendonça, Fabrizio Tagliavini, Isabel Santana, Chris Butler, Alex Gerhard, Adrian Danek, Johannes Levin, Markus Otto, Giovanni Frisoni, Roberta Ghidoni, Sandro Sorbi, Jonathan D Rohrer, Simon Ducharme, D. Louis Collins, Frontotemporal Lobar Degeneration Neuroimaging Initiative (FTLDNI), GENetic Frontotemporal dementia Initiative (GENFI)

**Affiliations:** McConnell Brain Imaging Centre, Montreal Neurological Institute, McGill University, Montreal, Quebec (QC), Canada. 3801, University, Montreal, Quebec, H3A 2B4.; Department of Psychiatry, McGill University Health Centre. 1025 Pine Avenue, Montreal, Quebec, Canada, H4A 1ª1.; Department of Neurology, Erasmus Medical Center, Rotterdam, The Netherlands; Centre for Neurodegenerative Disorders, Department of Clinical and Experimental Sciences, University of Brescia, Brescia, Italy; Alzheimer’s disease and Other Cognitive Disorders Unit, Neurology Service, Hospital Clínic, Institut d’Investigacións Biomèdiques August Pi I Sunyer, University of Barcelona, Barcelona, Spain; Cognitive Disorders Unit, Department of Neurology, Donostia University Hospital, San Sebastian, Gipuzkoa, Spain; Clinique Interdisciplinaire de Mémoire, Département des Sciences Neurologiques, CHU de Québec, and Faculté de Médecine, Université Laval, Quebec, Canada; Department of Geriatric Medicine, Karolinska University Hospital-Huddinge, Stockholm, Sweden; Department of Neurodegenerative Diseases, Hertie-Institute for Clinical Brain Research and Center of Neurology, University of Tübingen, Tübingen, Germany; Fondazione IRCCS Ca’ Granda Ospedale Maggiore Policlinico, Neurodegenerative Diseases Unit, Milan, Italy; LANE - Laboratory of Alzheimer's Neuroimaging and Epidemiology, IRCCS Istituto Centro San Giovanni di Dio Fatebenefratelli, Brescia, Italy; Department of Clinical Neurosciences, University of Cambridge, Cambridge, UK; Sunnybrook Health Sciences Centre, Sunnybrook Research Institute, University of Toronto, Toronto, Canada; Toronto Western Hospital, Tanz Centre for Research in Neurodegenerative Disease, Toronto, Ontario, Canada; Department of Clinical Neurological Sciences, University of Western Ontario, London, ON, Canada; Laboratory for Cognitive Neurology, Department of Neurosciences, KU Leuven, Leuven, Belgium; Faculty of Medicine, University of Lisbon, Lisbon, Portugal; Fondazione Istituto di Ricovero e Cura a Carattere Scientifico Istituto Neurologico Carlo Besta, Milan, Italy; Neurology Department, Centro Hospitalar e Universitário de Coimbra, Coimbra, Portugal; Department of Clinical Neurology, University of Oxford, Oxford, UK; Institute of Brain, Behaviour and Mental Health, The University of Manchester, Withington, Manchester, UK; Neurologische Klinik und Poliklinik, Ludwig-Maximilians-Universität, Munich, German Center for Neurodegenerative Diseases (DZNE), Munich, Germany; Department of Neurology, University Hospital Ulm, Ulm, Germany; Memory Clinic and LANVIE-Laboratory of Neuroimaging of Aging, University Hospitals and University of Geneva, Geneva, Switzerland; Molecular Markers Laboratory, IRCCS Istituto Centro San Giovanni di Dio Fatebenefratelli, Brescia, Italy; Department of Neuroscience, Psychology, Drug Research and Child Health, University of Florence, Florence, Italy; Department of Neuroscience, Psychology, Drug Research, and Child Health, University of Florence, Florence, Italy; Department of Neurodegenerative Disease, Dementia Research Centre, UCL Institute of Neurology, Queen Square, London, UK

**Keywords:** Frontotemporal dementia, Magnetic resonance, Deformation-based morphometry, Classification, Machine learning

## Abstract

**INTRODUCTION:** Brain structural imaging is paramount for the diagnosis of behavioral variant of frontotemporal dementia (bvFTD), but it has low sensitivity leading to erroneous or late diagnosis.

**METHODS:** A total of 515 subjects from two different bvFTD databases (training and validation cohorts) were included to perform voxel-wise deformation-based morphometry analysis to identify regions with significant differences between bvFTD and controls. A random forest classifier was used to individually predict bvFTD from morphometric differences in isolation and together with bedside cognitive scores.

**RESULTS:** Average ten-fold cross-validation accuracy was 89% (82% sensitivity, 93% specificity) using only MRI and 94% (89% sensitivity, 98% specificity) with the addition of semantic fluency. In a separate validation cohort of genetically confirmed bvFTD, accuracy was 88% (81% sensitivity, 92% specificity) with MRI and 91% (79% sensitivity, 96% specificity) with added cognitive scores.

**DISCUSSION:** The random forest classifier developed can accurately predict bvFTD at the individual subject level.

## 1. Introduction

The heterogeneity of frontotemporal dementia (FTD) is frequently considered a hallmark of the disease with significant variations in heritability, pathology and clinical presentations (1). First, although most cases of FTD are sporadic, there is a positive familiar history in 30-50% of the cases, and 10-30% are caused by an autosomal dominant mutation (most commonly progranulin-*GRN*-, microtubule-associated protein Tau-*MAPT*- and chromosome 9 open reading frame 72 - *C9orf72*-) (2, 3). Second, in terms of the underlying pathology, there are three main groups according to the major protein involved, all of which are characterized by selective degeneration of the frontal and temporal lobes: Tau, transactive response DNA-binding protein of 43 kDa - TDP-43-, and the tumor associated protein fused in sarcoma-FUS-.(4, 5)

Finally, regarding its clinical presentation, FTD is divided into three major clinical syndromes: the behavioral variant (bvFTD) and two language variants (Semantic Dementia and Non-Fluent Primary Progressive Aphasia). As the initially focal brain involvement spreads to involve larger regions in the frontal and temporal lobes, the symptoms of the three variants described can converge (5). Yet, there is also a considerable overlap between FTD and other neurodegenerative diseases; i.e. some patients with FTD may develop motor neuron disease, or symptoms of a Parkinsonian disorder (6).

Due to the aforementioned heterogeneity in pathology and heritability, as well as the syndromic overlap with psychiatric disorders and other dementias, a confirmed bvFTD diagnosis is often difficult to achieve in the absence of a dominant genetic mutation. Indeed, although brain imaging with magnetic resonance imaging (MRI) is paramount to increase the level of diagnostic confidence, it lacks sensitivity, particularly in the initial stages of the disease, leading to erroneous or late diagnosis (7, 8).

It has been recognized that the pattern of atrophy in sporadic bvFTD differed but also shared some similarities from that observed in mutation carriers. While frontal-particularly the anterior cingulate gyrus- and anterior temporal atrophy was associated to sporadic bvFTD, genetic variants showed distinct but overlapping patterns of atrophy (9–11). In *MAPT* mutation carriers, atrophy predominantly affects the anterior and medial temporal lobes, orbitofrontal lobe and insula; in *GRN* mutation carriers atrophy in the dorsolateral and ventromedial prefrontal, superolateral temporal and lateral parietal lobes as well as the anterior cingulate, insula, precuneus and striatum has been described; and *C9orf72* carriers showed relatively widespread cortical atrophy including posterior areas, and with particular involvement of the thalamus and superoposterior cerebellum (11).

Lately, machine learning techniques have been applied to distinguish between bvFTD and Cognitively Normal Subjects (CNCs), Alzheimer Disease or other psychiatric and neurologic disorders on an individual level using MRI based features (12–20). Studies vary greatly on the subjects included and the methodology. In the present study, we developed a Random Forest classifier (21) using features derived from Deformation Based Morphometry (DBM) maps to identify bvFTD subjects from CNCs. To ensure the generalizability of the results, the machine learning model was trained on a mainly sporadic cohort and tested in a held-out population of genetic bvFTD, therefore relying on one of the gold standards for bvFTD diagnosis (i.e., definite bvFTD)(7).

## 2. Materials and Methods

### 2.1. Participants

A total of 515 subjects were examined in this study. The first cohort (‘training cohort’) included bvFTD patients and CNCs from the Frontotemporal Lobar Degeneration Neuroimaging Initiative (FTLDNI) database who had T1-weighted (T1w) MRI scans matching with each clinical visit. Data was accessed and downloaded through the LONI platform in August 2018. The inclusion criteria for bvFTD patients was a diagnosis of possible or probable bvFTD according to the FTD consortium criteria (7), resulting in 70 patients with bvFTD and 123 CNCs in our study. The FTLDNI was funded through the National Institute of Aging and started in 2010. The primary goals of FTLDNI are to identify neuroimaging modalities and methods of analysis for tracking frontotemporal lobar degeneration (FTLD) and to assess the value of imaging versus other biomarkers in diagnostic roles. The project is the result of collaborative efforts at three sites in North America. For up-to-date information on participation and protocol, please visit: http://4rtni-ftldni.ini.usc.edu/

The second cohort (‘validation cohort’) included bvFTD patients and CNCs from the third data freeze (12/2017) of the Genetic Frontotemporal Dementia Initiative 2 (GENFI2-http://genfi.org.uk/) (22), which consists of 23 centres in the UK, Italy, The Netherlands, Sweden, Portugal and Canada. GENFI2 participants included known symptomatic carriers of a pathogenic mutation in *C9orf72*, *GRN* or *MAPT* and their first-degree relatives who are at risk of carrying a mutation, but who did not show any symptoms (i.e., presymptomatic). Non-carriers were first-degree relatives of symptomatic carriers who did not carry the mutation. The inclusion and exclusion criteria are described in detail elsewhere (22). Since the aim of the present study was to differentiate bvFTD patients from CNCs, presymptomatic carriers and symptomatic carriers whose clinical diagnosis was other than bvFTD were excluded. Non-carriers were considered as CNCs for the purpose of this study. This validation cohort contained 75 patients with bvFTD and 247 CNCs.

### 2.2. Clinical assessment

All FTLDNI subjects were regularly assessed every six-months for clinical characteristics (motor, non-motor and neuropsychological performance) by site investigators. Neuropsychological assessment included Mini Mental State Examination (MMSE), Montreal Cognitive Assessment (MOCA), Frontotemporal lobar degeneration clinical dementia rating (FTLD-CDR), Clinical Global Impression (CGI), verbal fluency, Frontotemporal dementia rating scale (FRS) amongst other cognitive and functional scores.

### 2.3. Image acquisition and preprocessing

For the FTLDNI training cohort, 3.0T MRIs were acquired at three sites. In all sites, a volumetric MPRAGE sequence was used to acquire T1w images of the entire brain. The acquisition parameters of the T1w images, using volumetric MPRAGE sequence, were RT/ET/IT: 2.3/3/900 ms, flip angle 9°, matrix 256×240, slice thickness 1mm, voxel size 1×1mm.

For the GENFI2 validation cohort, participants underwent volumetric T1w MRI at multiple centers, according to the GENFI imaging protocol. Sites used different types of scanners: Siemens Trio 3T, SiemensSkyra3T, Siemens1.5T, Phillips3T, General Electric (GE) 1.5T and GE 3T. Scan protocols were designed at the outset of the study to ensure adequate matching between the scanners and image quality control.

The T1w scans of the subjects were pre-processed through our longitudinal pipeline (23) that included image denoising (24), intensity non-uniformity correction (25), and image intensity normalization into range (0−100) using histogram matching. Each native T1w volume from each timepoint was linearly registered first to the subject-specific template which was then registered to the ICBM152 template (26, 27). All images were then non-linearly registered to the ICBM152 template using ANTs diffeomorphic registration pipeline (28). The images were visually assessed by two experienced raters to exclude cases with significant imaging artifacts (e.g. motion, incomplete field of view) or inaccurate linear/nonlinear registrations. This visual assessment was performed without any knowledge of the diagnosis. Out of 1724 scans, only 43 (2.5%, 36 scans in GENFI2, and 7 in FTLDNI) did not pass this visual quality control. For the purpose of this study, scans from subjects other than bvFTD or CNCs, or those that did not have a matching clinical visit were excluded from this analysis. This resulted in a total of 515 subjects that were included in this study.

### 2.4. Deformation based morphometry

DBM analysis was performed using Montreal Neurological Institute (MNI) MINC tools (23). The principle of DBM is to warp each individual scan to the template by introducing high-dimensional deformations (29, 30). Then, the morphological differences between the two are encoded in the deformations estimated for the warp. The local deformation obtained from the non-linear transformations was used as a measure of tissue expansion or atrophy by computing the determinant of the Jacobian at each voxel. Local contractions can be interpreted as shrinkage (e.g., tissue atrophy) and local expansions are often related to ventricular or sulci enlargement. DBM was used to assess both voxel-wise and atlas-based cross-sectional group related volumetric differences.

### 2.5. Classification bvFTD versus CNCs

To obtain a region of interest reflecting the difference between bvFTD and CNCs, a voxel-wise mixed effects model analysis was performed in the training dataset to assess the pattern of volumetric change according to diagnosis. The mixed effects model included age as a continuous fixed variable and diagnosis and sex as fixed categorical variables. Subject was included as a categorical random variable. The resulting maps were corrected for multiple comparisons using False Discovery Rate (FDR) with a *P value* < .05 threshold to identify regions associated with differences between bvFTD and CNCs; i.e. the diagnosis variable in the mixed effects model. A principal component analysis (PCA) was then performed on the DBM voxels within this region of interest. To avoid any leakage, only the training data was used for this PCA. Two sets of features were then used to train a random forests classifier (21) with 500 trees: 1) the first five principal components (PCs) as well as age and sex, and 2) the first five PCs, age, sex, and a neuropsychological score. The Semantic Fluency score (SF) was used as the cognitive score feature since it is a reliable simple bedside test associated with executive and language deficits in bvFTD (31) and was available for most of the subjects in both training and validation datasets. Executive deficits are considered a core characteristic of FTD, though in themselves, are insufficient to establish a diagnosis (32, 33). Classifications were run using DBM in isolation and DBM + SF. Ten-fold cross validation was used to assess the performance of the classifier within the training data.

To perform classification on the held-out GENFI2 validation dataset, the coefficients calculated based on the PCA on the training dataset were used to calculate the first five PCs features for the subjects from the validation dataset. Using the random forest classifier that was trained on FTLDNI, we then classified all the subjects from the validation dataset as either bvFTD or CNCs. A probability score was also obtained from the random forest classifier, indicating the likelihood of each observation belonging to the bvFTD class. The mixed effects modelling, PCA, and random forest classification were carried out using MATLAB (version R2017b).

### 2.6. Statistical analyses

All statistical analyses were also conducted using MATLAB (version R2017b). Two-sample t-Tests were conducted to examine demographic and clinical variables at baseline. Categorical variables were analysed using chi-square analyses. Results are expressed as mean ± standard deviation and median [interquartile range] as appropriate. Receiver Operator Characteristics (ROC) analysis was used to define sensitivity and specificity at different cut-points within the validation cohort. The optimal cut-point was estimated by the use of Youden index (J= Sensitivity+Specificity-1). Positive and negative likelihood ratios were also estimated for different cut points.

## 3. Results

### 3.1. Demographics

Table 1 shows the demographic and cognitive testing performances in bvFTD and CNCs. There was no difference in age between bvFTD patients and CNCs (62±6 and 63±6 years respectively, *P value* = 0.36), but there was a higher proportion of males in bvFTD patients than CNCs (67% vs 43%, *P value* = .001). As expected, bvFTD subjects showed greater cognitive and functional impairment: significant differences were found between the two cohorts in MMSE, FTLD-CDR, MOCA, letter fluency Z-score and semantic fluency Z-score (all *P value* < .001).

**Table 1.**
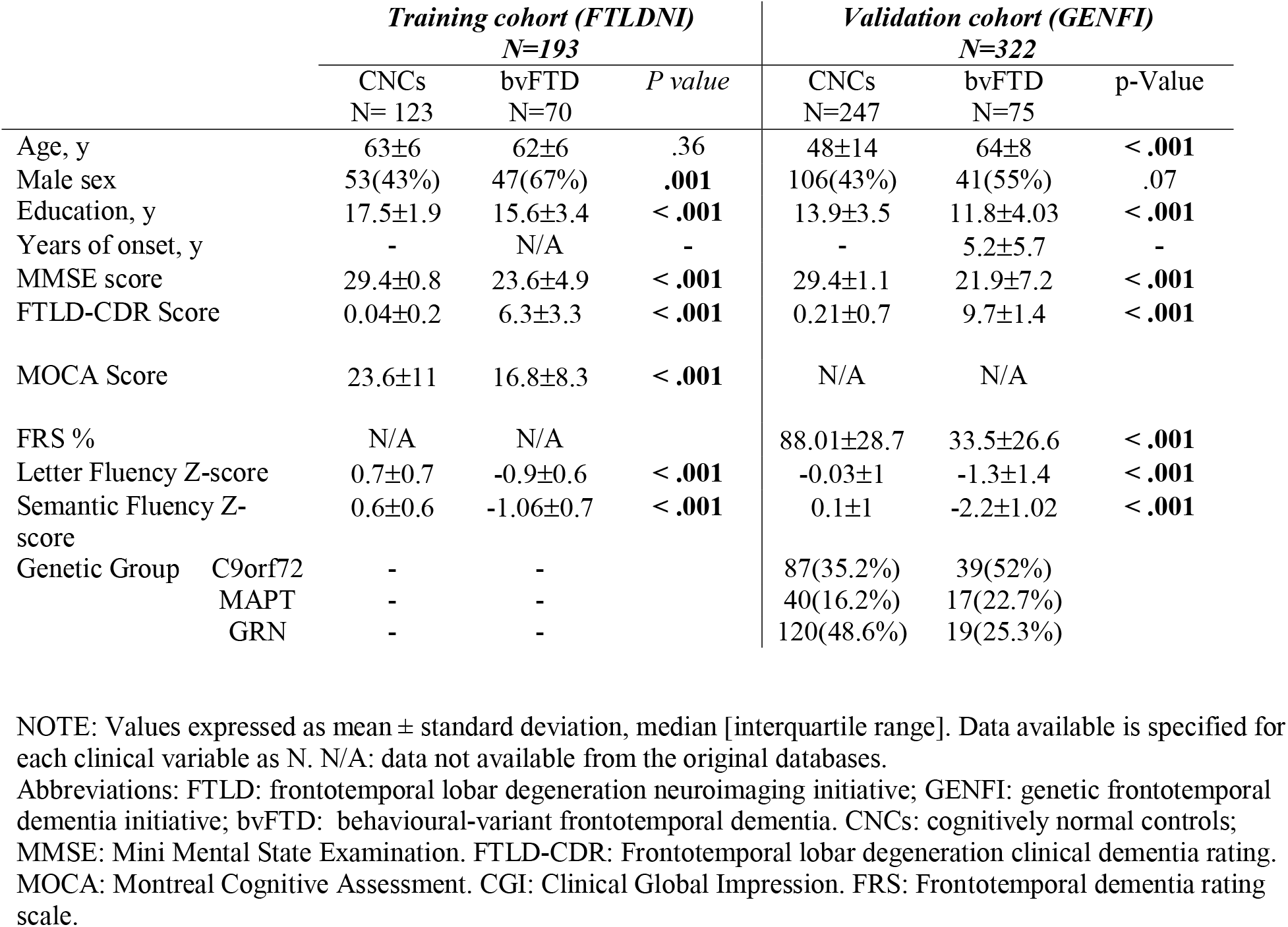
Demographic and clinical characteristics in bvFTD and healthy controls

Demographic differences and cognitive testing performances between patients and controls for the GENFI cohort are also shown in Table 1. Considering the CNCs from this dataset comes from non-carrier members of families at risk of genetic mutation related to FTD, they were, as expected, significantly younger than bvFTD subjects. The mean age was 48±14 years for CNCs and 64±8 years for bvFTD (*P value* < .001). The mean estimated disease duration for the bvFTD group was 5.2±5.7 years. Compared to non-carriers, bvFTD subjects showed greater cognitive and functional impairment. Significant differences were found between the two cohorts in MMSE, FTLD-CDR, MOCA, FRS, letter fluency Z-score and semantic fluency Z-score (*P value* < .001). Regarding the mutated gene, half of the bvFTD subjects carried a *C9orf72* mutation, while 22.7% and 25.3% belonged to the *MAPT* and *GRN* groups respectively.

### 3.2. Voxel-wise DBM group differences

Greater gray and white matter atrophy were found in the medial and inferior lateral portions of the frontal lobes as well as dorsolateral prefrontal cortex, insula, basal ganglia, medial and anterior temporal regions bilaterally and regions of brainstem and cerebellum in bvFTD. Correspondingly, volume increase was shown in the ventricles and sulci, being more evident in frontal horns and lateral sulcus (34).

### 3.3. Random forest classification

#### 3.3.1. Cross-validation results within the training cohort

The accuracy achieved for discrimination between bvFTD and CNCs using solely morphometric MRI features (DBM) was 89%, with a sensitivity of 82% and specificity of 93%. When adding one cognitive score (i.e., DBM+SF) the classifier accuracy reached 94%, with 89% sensitivity and 98% specificity.

#### 3.3.2. Classification within the validation cohort using solely DBM and DBM + SF

The application of the random forest classification model based on the training cohort to the validation cohort resulted in an accuracy of 88% when discriminating bvFTD patients from CNCs. Sensitivity and specificity were 81% and 92%, respectively using a probability score with an optimal cut point of 0.4 as threshold. This led to a positive likelihood ratio (LR+) of 10.13 and negative likelihood ratio (LR-) of 0.21.

The inclusion of semantic fluency in the classification model resulted in an accuracy of 91%, sensitivity of 79% and specificity of 96%; resulting in LR+ of 19.75 and LR- of 0.22. The ROC for DBM and DBM + SF classifiers are shown in Figure 1. Figure 2 shows the true positive rates for bvFTD and CNCs according the probability score for DBM (panel A) and DBM+SF (panel B). Table 2 shows the corresponding accuracy, sensitivity, specificity and likelihood ratios for the two models (DBM and DBM+SF) using different thresholds on the probability scores (e.g., for probability scores > 0.4).

**Table 2.** Classification performance using DBM and DBM + SF

|  | *Accuracy* | *Sensitivity* | *Specificity* | *LR+* | *LR-* |
| --- | --- | --- | --- | --- | --- |
| **DBM** |  |  |  |  |  |
| **Probability score** |  |  |  |  |  |
| **Threshold\*** |  |  |  |  |  |
| **0.60** | 0.90 | 0.53 | 0.98 | 26.50 | 0.48 |
| **0.55** | 0.91 | 0.64 | 0.97 | 21.33 | 0.37 |
| **0.50** | 0.90 | 0.71 | 0.96 | 17.75 | 0.30 |
| **0.45** | 0.89 | 0.77 | 0.93 | 11.00 | 0.25 |
| **0.40** | 0.88 | 0.81 | 0.92 | 10.13 | 0.21 |
| **0.35** | 0.86 | 0.81 | 0.91 | 9.00 | 0.21 |
| **0.30** | 0.84 | 0.83 | 0.88 | 6.92 | 0.19 |
| **0.25** | 0.81 | 0.87 | 0.87 | 6.69 | 0.15 |
| **0.20** | 0.78 | 0.89 | 0.83 | 5.24 | 0.13 |
| **0.15** | 0.70 | 0.91 | 0.74 | 3.50 | 0.12 |
| **0.10** | 0.60 | 0.95 | 0.64 | 2.64 | 0.08 |
| **DBM + SF** |  |  |  |  |  |
| **Probability score** |  |  |  |  |  |
| **Threshold\*** |  |  |  |  |  |
| **0.60** | 0.92 | 0.71 | 0.98 | 35.50 | 0.30 |
| **0.55** | 0.92 | 0.73 | 0.98 | 36.50 | 0.28 |
| **0.50** | 0.92 | 0.77 | 0.98 | 38.50 | 0.23 |
| **0.45** | 0.91 | 0.77 | 0.96 | 19.25 | 0.24 |
| **0.40** | 0.91 | 0.79 | 0.96 | 19.75 | 0.22 |
| **0.35** | 0.91 | 0.80 | 0.96 | 20.00 | 0.21 |
| **0.30** | 0.90 | 0.80 | 0.95 | 16.00 | 0.21 |
| **0.25** | 0.88 | 0.81 | 0.93 | 11.57 | 0.20 |
| **0.20** | 0.83 | 0.87 | 0.87 | 6.69 | 0.15 |
| **0.15** | 0.78 | 0.91 | 0.92 | 11.38 | 0.10 |
| **0.10** | 0.65 | 0.93 | 0.68 | 2.91 | 0.10 |

NOTE: \*Performances for each probability score threshold above which a subject is identified as bvFTD. Shaded rows correspond to the optimal cut-point estimated by Youden index.
Abbreviations: DBM: deformation-based morphometry; SF: semantic fluency score; LR+: positive likelihood ratio; LR-: negative likelihood ratio.

**Figure 1.**
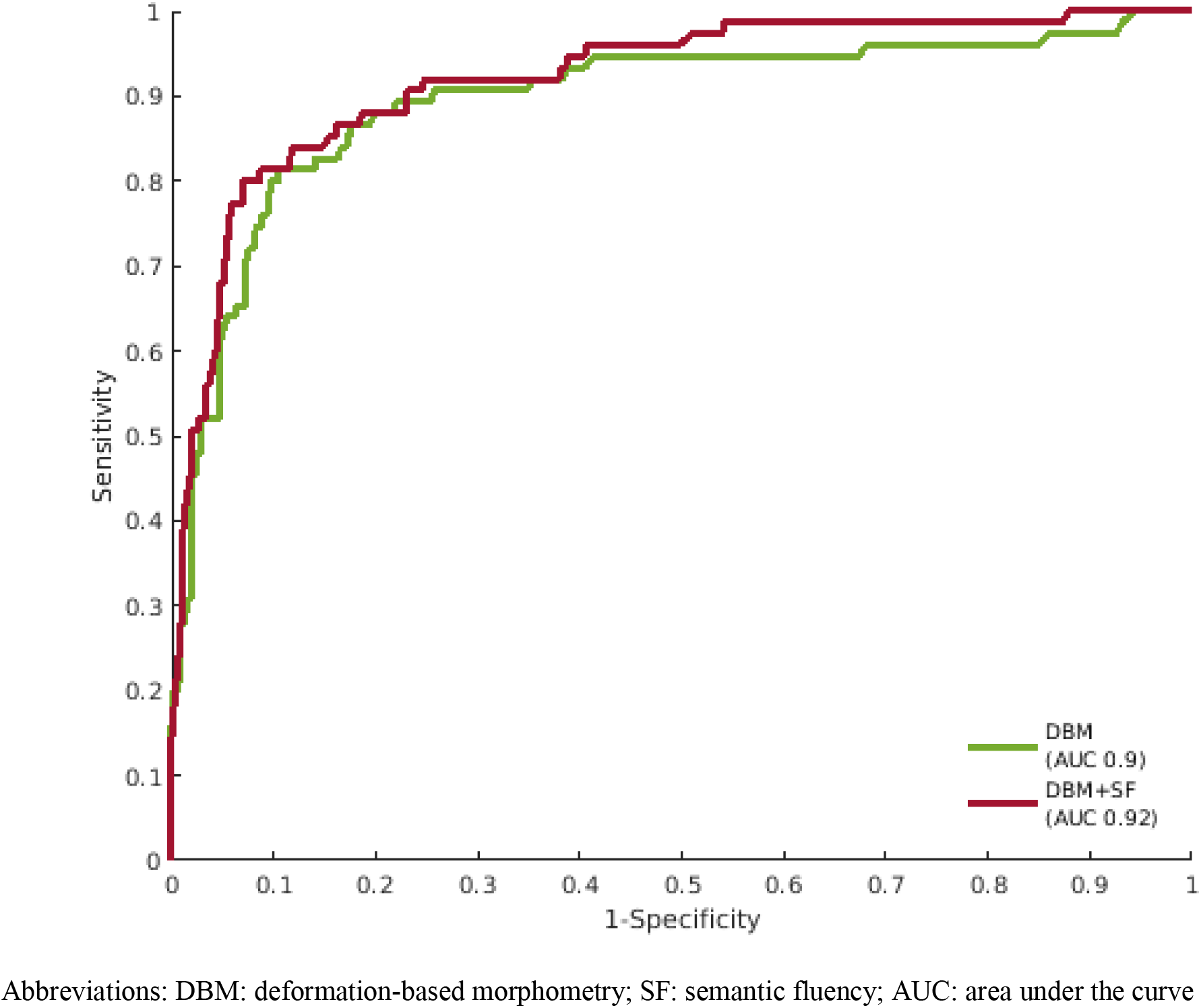
Receiver operating characteristic curves (ROC) for DBM and DBM+SF classifiers.

**Figure 2.**
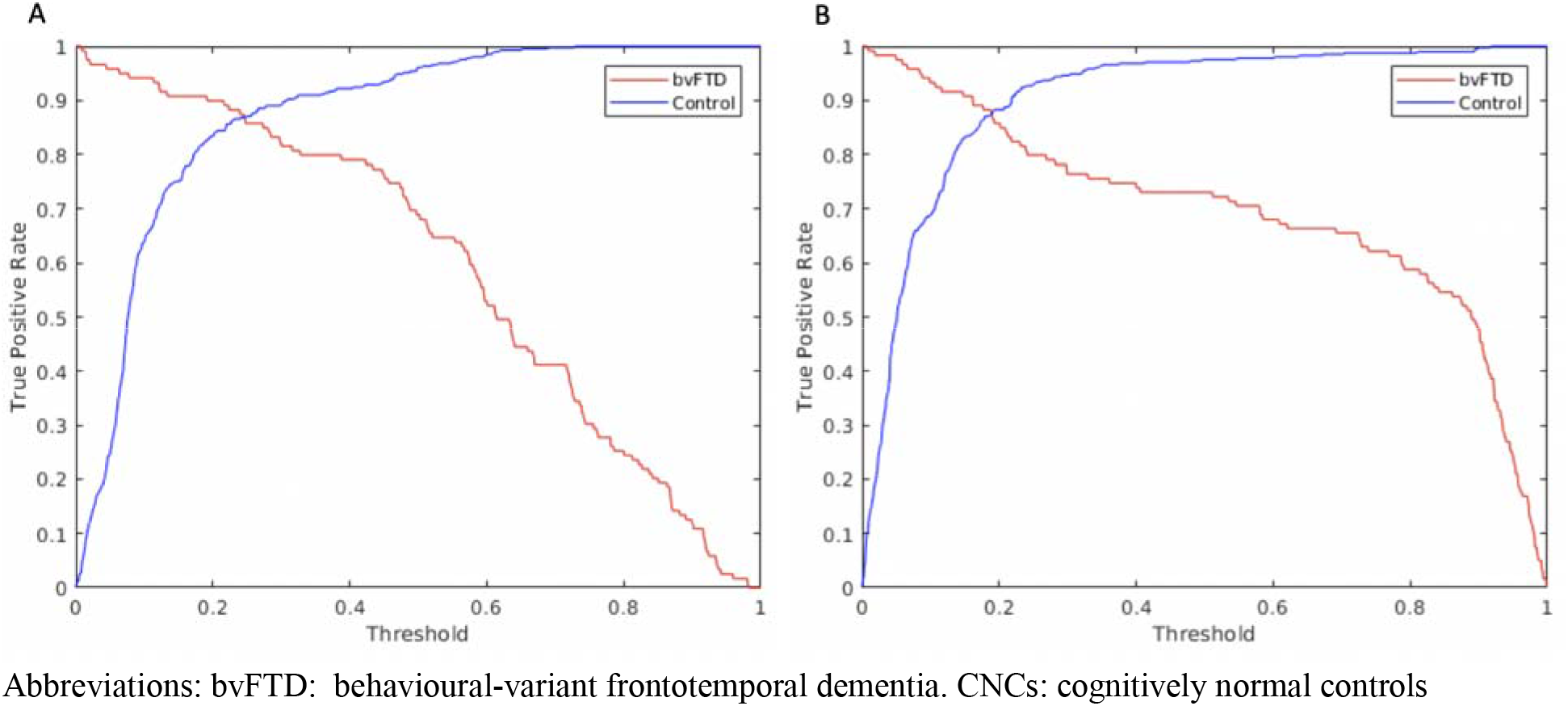
True positive rates for bvFTD and controls according to the probability score threshold for classification using DBM (panel A) or DBM + SF (panel B)

#### 3.3.3. False negative cases within the validation cohort

The classification using DBM resulted in 19% of false negatives. These subjects were significantly younger than the bvFTD subjects correctly classified (57±10 vs. 66±7 years respectively, *P value* < .001) and the estimated time from onset was also smaller (2±7 years vs 6±5 years; *P value* = .01). However, no significant differences were found in FTLD-CDR score between true positives and false negatives (*P value* = .07). The distribution of the genetic mutations did not show significant differences either between the false negatives and true positives. *GRN* corresponded to 22.7 % of all false negatives and 25.4% of all true positives (*P value* = .7); for *C9orf72* the distribution was 45.5% and 54.7% respectively (*P value* = .5) while for *MAPT* it was 31.8% of the false negatives and 18.9% of the correctly classified bvFTD (*P value* = .3)

#### 3.3.4. False positive cases within the validation cohort

Only 10 out of 247 CNCs (4%) were erroneously classified as bvFTD. These subjects were significantly older than the subjects accurately classified as healthy subjects (70±12 years vs. 47±13 years, respectively; *P value* < .001). No significant differences were found in the mean FTLD-CDR score (*P value* = .9). There was a small difference between the MMSE score for the false positives (28.27±2.2) and for true negatives (29.4±1; *P value* <.001).

#### 3.3.5. Defining strategic cut-points

Three cut-offs for both DBM and DBM+SF were defined by giving consideration to the sensitivity, specificity, positive and negative likelihood ratios of different points of the ROC: 1) the optimal cut-point according to Youden index; 2) a sensitive (i.e., “rule-out”) cut-point; and 3) a specific (i.e., “rule-in”) cut-point (Figure 3). The sensitivity, specificity, LR- and LR+ expressed in the figure were estimated for each of these defined cut-points (e.g., for probability score = 0.4)

**Figure 3.**
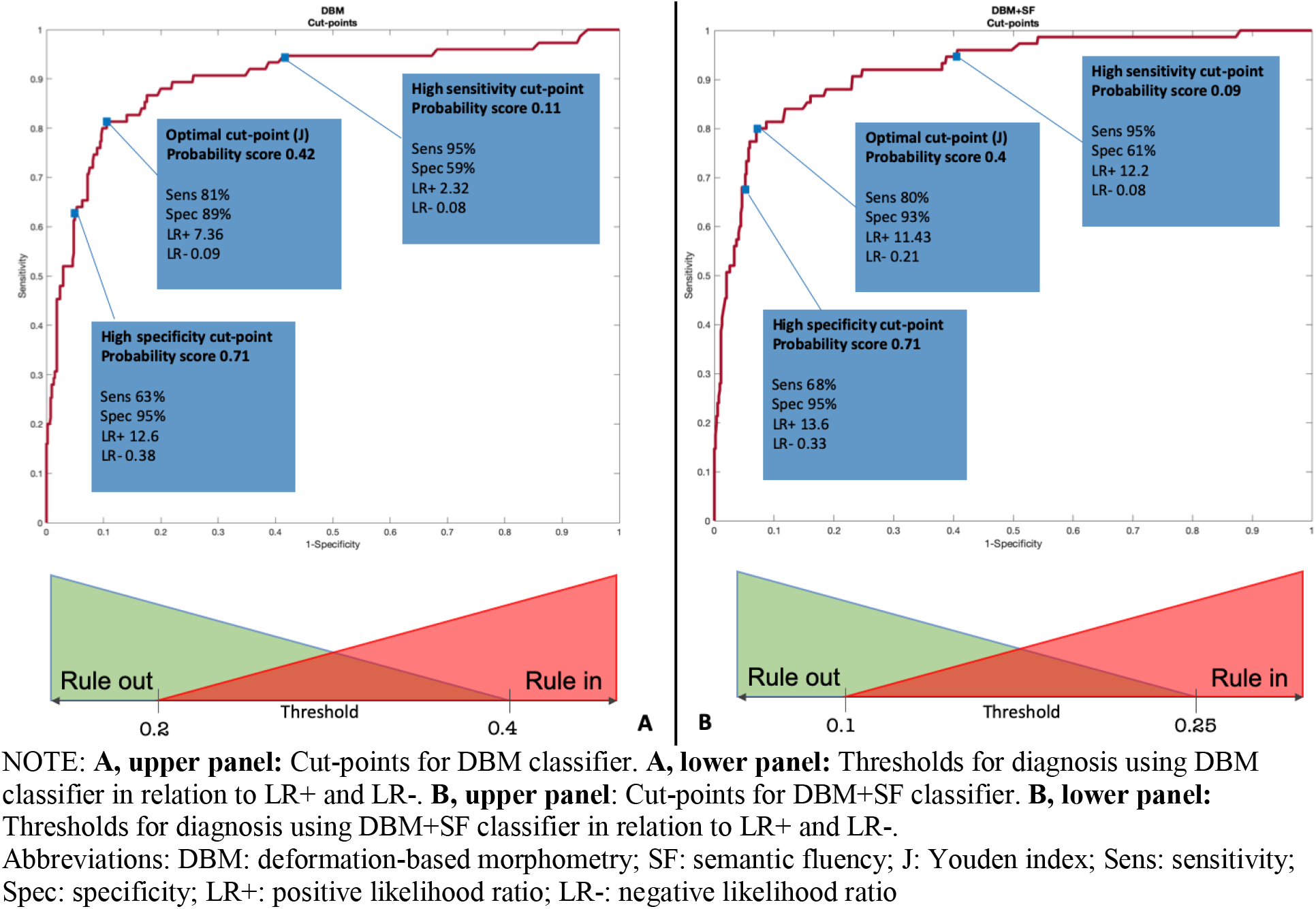
Strategic ROC cut-points.

Proposed thresholds for clinical decision-making for each classifier according to their likelihood ratios are proposed in Figure 3 (lower panels). A LR- <0.1 allows to reliably exclude (i.e., rule-out) bvFTD when the probability score is below 0.2 and 0.1 for DBM and DBM+SF, respectively. Probability scores over 0.4 for DBM and 0.25 for DBM+SF allow to confidently diagnose (i.e., rule-in) the disease with a LR+ >10. Corresponding likelihood ratios for different thresholds are shown in table 2.

## 4. Discussion

In the present study we built a random forest classifier using morphometric MRI features for the individual prediction of bvFTD. The main findings are: 1) our random forest algorithm yielded areas under the curve of 0.90 and 0.92 using DBM and DBM+SF, respectively, in the independent validation cohort of genetically confirmed bvFTD cases; 2) the inclusion of a simple cognitive score (SF) improved the accuracies and specificity regardless of the probability threshold chosen, while reducing the false negative rate for probability scores > 0.5; 3) we provide three cut-off values (a “statistically optimal” cut-point, a sensitive (“rule-out”) cut-point and a specific (“rule-in”) cut-point) for both DBM and DBM+SF classifiers; and 4) our results show good positive and negative likelihood ratios proving its reliability in ruling in and out the disease.

The likelihood ratio is the percentage of patients with a given test result divided by the percentage of controls with the same results. Meaning that ill people should be more likely to have an abnormal result of a given test than healthy individuals(35, 36). For DBM only classifier in the independent validation cohort, the optimal threshold yielded an area under the curve (AUC) of 0.9 with 81% sensitivity and 92% specificity leading to a positive LR+ of 10.13 and negative LR- of 0.21. Whereas, the AUC, sensitivity and specificity using the DBM+SF model were 0.92, 79% and 96%, respectively. These values result in LR+ of 19.75 and LR- of 0.22. To keep in mind, a test is moderately good at ruling in disease when LR+ is greater than 2 and very good at doing it when LR+ is greater than 10 (37). Furthermore, a test is moderately good at ruling out the disease with LR- below 0.5 and very good below 0.1. Hence, using the optimal thresholds, both models are very good at excluding non bvFTD subjects and moderately good at confirming the disease.

Our results show that the random forest classifier we developed in our training cohort can accurately predict bvFTD in individual subjects in a completely independent validation cohort coming from a different and independent database. Furthermore, the GENFI2 validation cohort includes bvFTD patients with a definite diagnosis (positive genetic mutation). Of note, our algorithm was able to accurately classify patients with genetic bvFTD even though they tend to have more atypical atrophy patterns(11). The performance of our classifier is superior than the performance reported in several articles that have analyzed the standard diagnostic methods currently used in the clinical practice. Within a pathology-confirmed cohort the sensitivity reported for the revised diagnostic criteria for bvFTD was 86% for possible diagnosis and 75% for probable bvFTD (with neuroimaging support) (7). However, these criteria reported a sensitivity of 85% and specificity of 27% for possible bvFTD diagnosis in a clinically relevant cohort of patients with mixed behavioral changes, reaching 82% specificity when adding a compatible MRI scan (38). Within a cohort with late onset behavioral disorders, 70% sensitivity and 93% specificity have been reported for structural MRI alone for bvFTD assessed by an experienced neuroradiologist (8). The latter results have comparable positive and negative likelihood ratios to ours, even though our method does not rely on the expertise of the radiological observer.

Previous studies classified bvFTD from a control group (12–16). The best AUC was reported by Raamana et al (AUC 0.938, 100% sensitivity and 88% specificity). However, the main limitation of that study is that the bvFTD diagnosis from the validation cohort was based on clinical criteria (39). Contrarily, the bvFTD subjects from our validation cohort are known carriers of a pathogenic mutation and have therefore, definite bvFTD diagnosis.

The performance of the classifier was tested on a held-out database which included multi-center and multi-scanner data from different scanner models of both 1.5T and 3T field strengths. This further reinforces the generalizability (i.e., external validity) of our results and ensures their applicability in a clinical scenario with different scanners, even with different magnetic field strengths. This certainly constitutes one of the main strengths of this study along with the fact that our performance was estimated using one of the gold standards for FTD diagnosis (i.e., definite FTD supported by the presence of a pathogenic mutation). Of significant value, our algorithm is based on standard structural T1w MRI and a simple cognitive test that are already routinely acquired clinically, making for strong translational potential. On the other hand, the main limitation is that these results are yet to be validated prospectively in a clinically representative cohort including patients with diverse primary psychiatric disorders (a common differential diagnosis from bvFTD) (40). The classification accuracy also remains to be demonstrated in cohorts with other types of dementias and cardiovascular comorbidities, as these were uncommon in our dataset and could have influenced our very high specificity. Finally, in our results the false negatives/positives were significantly younger/older than the subjects that were correctly classified. This is likely due to the fact that the age range for the validation dataset (GENFI: minimum age: 39, maximum age:79) was larger than the training set (FTLDNI, minimum age 46, maximum age 75). Subjects that were outside the operating range of the classifier were therefore more likely to be misclassified. Adding subjects with similar ages to the training dataset will likely improve the results. In addition, specifically for the false negative cases, although the difference did not reach statistical significance (p=0.07), the false negatives had lower FTLD-CDR scores than the true positive cases, implying an earlier stage of the disease. It is plausible that such subjects with milder symptoms were not as well represented in NIFD given the difficulty of diagnosing bvFTD in the very mild stages when there is no known genetic mutation.

To conclude, we propose an automatic method using structural MRI alone (already available and routinely used in the clinic) and including a simple cognitive test that could be administered by any physician in few minutes for reliable individual prediction of bvFTD at the individual subject level. If validated in a prospective study, this algorithm has the potential to improve diagnostic accuracy, particularly in setting without access to specialized FTD care.

## Abbreviations

FTD: frontotemporal dementia
GRN: progranulin
MAPT: microtubule-associated protein tau
C9orf72: chromosome 9 open reading frame 72
bvFTD: behavioural variant of frontotemporal dementia
MRI: magnetic resonance imaging
CNCs: cognitively normal controls
DBM: deformation-based morphometry
FTLDNI: frontotemporal lobar degeneration neuroimaging initiative
T1-w: T1 weighted
GENFI: Genetic frontotemporal dementia initiative
MMSE: Mini mental state examination
MOCA: Montreal cognitive assessment
FTLD-CDR: Frontotemporal lobar degeneration Clinical Dementia Rating score
CGI: Clinical global impression
FRS: Frontotemporal dementia rating scale
FDR: False Discovery Rate
PCA: Principal component analysis
PCs: Principal components
SF: Semantic fluency
ROC: Receiver operating characteristic curves
AUC: Area under the curve
LR+: positive likelihood ratio
LR−: negative likelihood ratio

## Acknowledgements

We would like to acknowledge funding from the Famille Louise & André Charron.

Data collection and sharing for this project was funded by the Frontotemporal Lobar Degeneration Neuroimaging Initiative (National Institutes of Health Grant R01 AG032306). The study is coordinated through the University of California, San Francisco, Memory and Aging Center. FTLDNI data are disseminated by the Laboratory for Neuro Imaging at the University of Southern California.

Brain scan acquisition at the McConnell Brain Imaging was supported by the Brain Canada Foundation with support from Health Canada and the Canada Foundation for Innovation (CFI Project 34874).

This work was supported by Italian Ministry of Health (CoEN015 and Ricerca Corrente).

## Declaration of interests

**Dr. Manera** reports no disclosures

**Dr. Dadar** reports no disclosures

**Dr. Collins** is co-founder of True Positive Medical Devices.

**Dr. Ducharme** receives salary funding from the Fonds de Recherche du Québec - Santé. Dr. Ducharme is the co-founder of Arctic Fox AI.

**Dr. Ghidoni** reports no disclosures

## Appendix List of other GENFI consortium members

Sónia Afonso - Instituto Ciencias Nucleares Aplicadas a Saude, Universidade de Coimbra, Coimbra, Portugal Maria Rosario Almeida - Centre of Neurosciences and Cell Biology, Universidade de Coimbra, Coimbra, Portugal Sarah Anderl-Straub – Department of Neurology, Ulm University, Ulm, Germany

Christin Andersson - Department of Clinical Neuroscience, Karolinska Institutet, Stockholm, Sweden

Anna Antonell - Alzheimer’s disease and other cognitive disorders unit, Neurology Department, Hospital Clinic, Institut d’Investigacions Biomèdiques, Barcelona, Spain

Silvana Archetti - Biotechnology Laboratory, Department of Diagnostics, ASST Brescia Hospital, Brescia, Italy Andrea Arighi - Fondazione IRCSS Ca’ Granda, Ospedale Maggiore Policlinico, Neurodegenerative Diseases Unit, Milan, Italy

Mircea Balasa - Alzheimer’s disease and other cognitive disorders unit, Neurology Department, Hospital Clinic, Institut d’Investigacions Biomèdiques, Barcelona, Spain

Myriam Barandiaran - Neuroscience Area, Biodonostia Health Research Institute, Paseo Dr Begiristain sn, CP 20014, San Sebastian, Gipuzkoa, Spain

Nuria Bargalló - Radiology Department, Image Diagnosis Center, Hospital Clínic and Magnetic Resonance Image core facility, IDIBAPS, Barcelona, Spain

Robart Bartha - Department of Medical Biophysics, Robarts Research Institute, University of Western Ontario, London, Ontario, Canada

Benjamin Bender - Department of Diagnostic and Interventional Neuroradiology, University of Tuebingen, Tuebingen, Germany

Alberto Benussi - Centre for Neurodegenerative Disorders, Department of Clinical and Experimental Sciences, University of Brescia, Italy

Luisa Benussi - Istituto di Ricovero e Cura a Carattere Scientifico Istituto Centro San Giovanni di Dio Fatebenefratelli, Brescia, Italy

Valentina Bessi - Department of Neuroscience, Psychology, Drug Research, and Child Health, University of Florence, Florence, Italy

Giuliano Binetti - Istituto di Ricovero e Cura a Carattere Scientifico Istituto Centro San Giovanni di Dio Fatebenefratelli, Brescia, Italy

Sandra Black - LC Campbell Cognitive Neurology Research Unit, Sunnybrook Research Institute, Toronto, Canada Martina Bocchetta – Dementia Research Centre, Department of Neurodegenerative Disease, UCL Institute of Neurology, Queen Square London, UK

Sergi Borrego-Ecija - Alzheimer’s disease and other cognitive disorders unit, Neurology Department, Hospital Clinic, Institut d’Investigacions Biomèdiques, Barcelona, Spain

Jose Bras – Dementia Research Institute, Department of Neurodegenerative Disease, UCL Institute of Neurology, Queen Square, London, UK

Rose Bruffaerts - Laboratory for Cognitive Neurology, Department of Neurosciences, KU Leuven, Leuven, Belgium Paola Caroppo - Fondazione Istituto di Ricovero e Cura a Carattere Scientifico Istituto Neurologico Carlo Besta, Milan, Italy

David Cash – Dementia Research Centre, Department of Neurodegenerative Disease, UCL Institute of Neurology, Queen Square, London, UK

Miguel Castelo-Branco - Neurology Department, Centro Hospitalar e Universitário de Coimbra, Instituto de Ciências Nucleares Aplicadas à Saúde (ICNAS), Coimbra, Portugal

Rhian Convery – Dementia Research Centre, Department of Neurodegenerative Disease, UCL Institute of Neurology, Queen Square, London, UK

Thomas Cope – Department of Clinical Neuroscience, University of Cambridge, Cambridge, UK Maura Cosseddu – Neurology, ASST Brescia Hospital, Brescia, Italy

María de Arriba - Neuroscience Area, Biodonostia Health Research Institute, Paseo Dr Begiristain sn, CP 20014, San Sebastian, Gipuzkoa, Spain

Giuseppe Di Fede - Fondazione Istituto di Ricovero e Cura a Carattere Scientifico Istituto Neurologico Carlo Besta, Milan, Italy

Zigor Díaz - CITA Alzheimer, San Sebastian, Spain

Diana Duro - Faculty of Medicine, Universidade de Coimbra, Coimbra, Portugal Chiara Fenoglio - University of Milan, Centro Dino Ferrari, Milan, Italy

Camilla Ferrari - Department of Neuroscience, Psychology, Drug Research, and Child Health, University of Florence, Florence, Italy

Carlos Ferreira - Instituto Ciências Nucleares Aplicadas à Saúde, Universidade de Coimbra, Coimbra, Portugal Catarina B. Ferreira - Faculty of Medicine, University of Lisbon, Lisbon, Portugal

Toby Flanagan – Faculty of Biology, Medicine and Health, Division of Neuroscience and Experimental Psychology, University of Manchester, Manchester, UK

Nick Fox – Dementia Research Centre, Department of Neurodegenerative Disease, UCL Institute of Neurology, Queen Square, London, UK

Morris Freedman - Division of Neurology, Baycrest Centre for Geriatric Care, University of Toronto, Toronto, Canada

Giorgio Fumagalli - Fondazione IRCSS Ca’ Granda, Ospedale Maggiore Policlinico, Neurodegenerative Diseases Unit, Milan, Italy; Department of Neuroscience, Psychology, Drug Research and Child Health, University of Florence, Florence, Italy

Alazne Gabilondo - Neuroscience Area, Biodonostia Health Research Institute, Paseo Dr Begiristain sn, CP 20014, San Sebastian, Gipuzkoa, Spain

Roberto Gasparotti - Neuroradiology Unit, University of Brescia, Brescia, Italy

Serge Gauthier - Department of Neurology and Neurosurgery, McGill University, Montreal, Québec, Canada Stefano Gazzina - Neurology, ASST Brescia Hospital, Brescia, Italy

Giorgio Giaccone - Fondazione Istituto di Ricovero e Cura a Carattere Scientifico Istituto Neurologico Carlo Besta, Milan, Italy

Ana Gorostidi - Neuroscience Area, Biodonostia Health Research Institute, Paseo Dr Begiristain sn, CP 20014, San Sebastian, Gipuzkoa, Spain

Caroline Greaves – Dementia Research Centre, Department of Neurodegenerative Disease, UCL Institute of Neurology, Queen Square London, UK

Rita Guerreiro – Dementia Research Institute, Department of Neurodegenerative Disease, UCL Institute of Neurology, London, UK

Carolin Heller – Dementia Research Centre, Department of Neurodegenerative Disease, UCL Institute of Neurology, Queen Square, London, UK

Tobias Hoegen - Department of Neurology, Ludwig-Maximilians-University of Munich, Munich, Germany

Begoña Indakoetxea - Cognitive Disorders Unit, Department of Neurology, Donostia University Hospital, Paseo Dr Begiristain sn, CP 20014, San Sebastian, Gipuzkoa, Spain

Vesna Jelic - Division of Clinical Geriatrics, Karolinska Institutet, Stockholm, Sweden

Lize Jiskoot - Department of Neurology, Erasmus Medical Center, Rotterdam, The Netherlands

Hans-Otto Karnath - Section of Neuropsychology, Department of Cognitive Neurology, Center for Neurology & Hertie-Institute for Clinical Brain Research, Tübingen, Germany

Ron Keren - University Health Network Memory Clinic, Toronto Western Hospital, Toronto, Canada

Maria João Leitão - Centre of Neurosciences and Cell Biology, Universidade de Coimbra, Coimbra, Portugal

Albert Lladó - Alzheimer’s disease and other cognitive disorders unit, Neurology Department, Hospital Clinic, Institut d’Investigacions Biomèdiques, Barcelona, Spain

Gemma Lombardi - Department of Neuroscience, Psychology, Drug Research and Child Health, University of Florence, Florence, Italy

Sandra Loosli - Department of Neurology, Ludwig-Maximilians-University of Munich, Munich, Germany

Carolina Maruta - Lisbon Faculty of Medicine, Language Research Laboratory, Lisbon, Portugal

Simon Mead - MRC Prion Unit, Department of Neurodegenerative Disease, UCL Institute of Neurology, Queen Square, London, UK

Lieke Meeter - Department of Neurology, Erasmus Medical Center, Rotterdam, Netherlands Gabriel Miltenberger - Faculty of Medicine, University of Lisbon, Lisbon, Portugal

Rick van Minkelen - Department of Clinical Genetics, Erasmus Medical Center, Rotterdam, The Netherlands

Sara Mitchell - LC Campbell Cognitive Neurology Research Unit, Sunnybrook Research Institute, Toronto, Canada Katrina M Moore – Dementia Research Centre, Department of Neurodegenerative Disease, UCL Institute of Neurology, Queen Square, London, UK

Benedetta Nacmias - Department of Neuroscience, Psychology, Drug Research and Child Health, University of Florence, Florence, Italy

Mollie Neason - Dementia Research Centre, Department of Neurodegenerative Disease, UCL Institute of Neurology, Queen Square, London, UK

Jennifer Nicholas – Department of Medical Statistics, London School of Hygiene and Tropical Medicine, London, UK

Linn Öijerstedt - Department of Geriatric Medicine, Karolinska Institutet, Stockholm, Sweden

Jaume Olives - Alzheimer’s disease and other cognitive disorders unit, Neurology Department, Hospital Clinic, Institut d’Investigacions Biomèdiques, Barcelona, Spain

Sebastien Ourselin - School of Biomedical Engineering & Imaging Sciences, King's College London, London, UK Alessandro Padovani - Centre for Neurodegenerative Disorders, Department of Clinical and Experimental Sciences, University of Brescia, Italy

Jessica Panman – Department of Neurology, Erasmus Medical Center, Rotterdam, The Netherlands Janne Papma - Department of Neurology, Erasmus Medical Center, Rotterdam, The Netherlands

Georgia Peakman - Department of Neurodegenerative Disease, Dementia Research Centre, UCL Institute of Neurology, Queen Square, London, UK

Irene Piaceri - Department of Neuroscience, Psychology, Drug Research and Child Health, University of Florence, Florence

Michela Pievani - Istituto di Ricovero e Cura a Carattere Scientifico Istituto Centro San Giovanni di Dio Fatebenefratelli, Brescia, Italy

Yolande Pijnenburg - VUMC, Amsterdam, The Netherlands

Cristina Polito - Department of Biomedical, Experimental and Clinical Sciences “Mario Serio”, Nuclear Medicine Unit, University of Florence, Florence, Italy

Enrico Premi - Stroke Unit, ASST Brescia Hospital, Brescia, Italy

Sara Prioni - Fondazione Istituto di Ricovero e Cura a Carattere Scientifico Istituto Neurologico Carlo Besta, Milan, Italy

Catharina Prix - Department of Neurology, Ludwig-Maximilians-University Munich, Germany Rosa Rademakers - Department of Neurosciences, Mayo Clinic, Jacksonville, Florida, USA

Veronica Redaelli - Fondazione Istituto di Ricovero e Cura a Carattere Scientifico Istituto Neurologico Carlo Besta, Milan, Italy

Tim Rittman – Department of Clinical Neurosciences, University of Cambridge, Cambridge, UK

Ekaterina Rogaeva - Tanz Centre for Research in Neurodegenerative Diseases, University of Toronto, Toronto, Canada

Pedro Rosa-Neto - Translational Neuroimaging Laboratory, McGill University Montreal, Québec, Canada Giacomina Rossi - Fondazione Istituto di Ricovero e Cura a Carattere Scientifico Istituto Neurologico Carlo Besta, Milan, Italy

Martin Rossor – Dementia Research Centre, Department of Neurodegenerative Disease, UCL Institute of Neurology, Queen Square, London, UK

Beatriz Santiago - Neurology Department, Centro Hospitalar e Universitário de Coimbra, Coimbra, Portugal

Elio Scarpini - University of Milan, Centro Dino Ferrari, Milan, Italy; Fondazione IRCSS Ca’ Granda, Ospedale Maggiore Policlinico, Neurodegenerative Diseases Unit, Milan, Italy

Sonja Schönecker - Neurologische Klinik, Ludwig-Maximilians-Universität München, Munich, Germany Elisa Semler – Department of Neurology, Ulm University, Ulm, Germany

Rachelle Shafei – Dementia Research Centre, Department of Neurodegenerative Disease, UCL Institute of Neurology, Queen Square, London, UK

Christen Shoesmith - Department of Clinical Neurological Sciences, University of Western Ontario, London, Ontario, Canada

Miguel Tábuas-Pereira - Centre of Neurosciences and Cell Biology, Universidade de Coimbra, Coimbra, Portugal Mikel Tainta - Neuroscience Area, Biodonostia Health Research Institute, Paseo Dr Begiristain sn, CP 20014, San Sebastian, Gipuzkoa, Spain

Ricardo Taipa - Neuropathology Unit and Department of Neurology, Centro Hospitalar do Porto - Hospital de Santo António, Oporto, Portugal

David Tang-Wai - University Health Network Memory Clinic, Toronto Western Hospital, Toronto, Canada David L Thomas - Neuroradiological Academic Unit, UCL Institute of Neurology, London, UK

Hakan Thonberg - Center for Alzheimer Research, Division of Neurogeriatrics, Karolinska Institutet, Stockholm, Sweden

Carolyn Timberlake - Department of Clinical Neurosciences, University of Cambridge, Cambridge, UK

Pietro Tiraboschi - Fondazione Istituto di Ricovero e Cura a Carattere Scientifico Istituto Neurologico Carlo Besta, Milano, Italy

Emily Todd - Department of Neurodegenerative Disease, Dementia Research Centre, UCL Institute of Neurology, Queen Square, London, UK

Philip Vandamme - Neurology Service, University Hospitals Leuven, Belgium; Laboratory for Neurobiology, VIB-KU Leuven Centre for Brain Research, Leuven, Belgium

Mathieu Vandenbulcke - Geriatric Psychiatry Service, University Hospitals Leuven, Belgium; Neuropsychiatry, Department of Neurosciences, KU Leuven, Leuven, Belgium

Michele Veldsman - Nuffield Department of Clinical Neurosciences, Medical Sciences Division, University of Oxford, UK

Ana Verdelho - Department of Neurosciences, Santa Maria Hospital, University of Lisbon, Portugal Jorge Villanua - OSATEK Unidad de Donostia, San Sebastian, Gipuzkoa, Spain

Jason Warren – Dementia Research Centre, Department of Neurodegenerative Disease, UCL Institute of Neurology, Queen Square, London, UK

Carlo Wilke - Hertie Institute for Clinical Brain Research, University of Tuebingen, Tuebingen, Germany Ione Woollacott – Dementia Research Centre, Department of Neurodegenerative Disease, UCL Institute of Neurology, Queen Square, London, UK

Elisabeth Wlasich - Neurologische Klinik, Ludwig-Maximilians-Universität München, Munich, Germany Henrik Zetterberg - Department of Neurodegenerative Disease, UCL Institute of Neurology, London, UK

Miren Zulaica - Neuroscience Area, Biodonostia Health Research Institute, Paseo Dr Begiristain sn, CP 20014, San Sebastian, Gipuzkoa, Spain.

